# Experiments on localization accuracy with non-individual and individual HRTFs comparing static and dynamic reproduction methods

**DOI:** 10.1101/2020.03.31.011650

**Authors:** Josefa Oberem, Jan-Gerrit Richter, Dorothea Setzer, Julia Seibold, Iring Koch, Janina Fels

## Abstract

Binaural reproduction can be used in listening experiments under real-life conditions to achieve a high realism and good reproducibility. In recent years a clear trend to more individual reproduction can be observed as the ability to measure individual head-related-transfer-functions (HRTFs) is becoming more widespread. The question of the accuracy and reproduction methods needed for a realistic playback however has not been sufficiently answered. To evaluate an appropriate approach for binaural reproduction via headphones different head-related-transfer-functions (HRTFs) and reproduction methods were compared in this paper. In a listening test eleven explicitly trained participants were asked to localize eleven sound sources positioned in the right hemisphere using the proximal pointing method. Binaural stimuli based on individually measured HRTFs were compared to those of an artificial head in a static reproduction of stimuli and in three dynamic reproduction methods of different resolutions (5°, 2.5° and 1°). Unsigned errors in azimuth and elevation as well as front-back-confusions and in-head-localization were observed. Dynamic reproduction of any resolution applied turned out fundamental for a reduction of undesired front-back-confusions and in-head-localization. Individually measured HRTFs showed a smaller effect on localization accuracy compared to the influence of dynamic sound reproduction. They were mainly observed to reduce the front-back-confusion rate.

## 1 Introduction

Parts of this study were presented at the conference DAGA, Munich, Germany, 2018 [1] and are published in the PhD-thesis [2].

The progress of individual binaural reproduction of recent years has brought the capability to study complex cognitive processes which require real-life conditions and high level of plausibility under a controlled and reproduce-able environment. The accuracy and plausibility of these binaural reproductions is highly dependent on several factors. These factors range from, for example, the set of head-related-transfer-functions (HRTFs) used (e.g. individuality, resolution, section) as well as the type of reproduction (e.g. dynamic, static). The importance of each of these components individually has been discussed for several years. The benefit of the use of individual HRTFs for localization tasks has been discussed for several decades with some of the earliest work by Wightman, Kistler and Wenzel [3], [4]. A gain from small head movements was described first in Wallach [5] and could be reproduced several times [6]–[8].

However, no study has systematically looked at the influence of the *resolution* of high resolution individual HRTFs and tried to quantify the influence of *both* high resolution individual HRTFs as well as dynamic binaural reproduction. Consequently, when dealing with a reproduction problem that requires real-life conditions, one must assume that all factors will benefit equally and are crucially important.

As taking all these factors into account is a time-consuming and expensive procedure the goal of this publication is to find a HRTF dataset which, combined with a reproduction method (static / dynamic), will result in a performance comparable to performance achieved with real sources. To this end localization experiments were conducted since localization performance delivers a rather general measure to indicate the quality of binaural auditory displays, not limited by any paradigm conditions. In the experiment individually measured HRTFs were compared to those of an artificial head in a static reproduction of stimuli and in three dynamic reproduction methods of different resolutions (5°, 2.5° and 1°).

## 2 State of the Art

Several studies investigated localization performance for real sources in comparison with virtual sound sources using a binaural headphone reproduction. There was a broad agreement that results nearly as accurate as results for real sources could also be achieved by using individual HRTFs for sound reproduction [3], [9]–[11]. Headphone reproduction, however, featured a generally higher rate of front-back-confusions (e.g. [3]). Investigations on experiments using individual and non-individual HRTF measurements demonstrated the necessity to use individual HRTFs in order to attain results nearly equivalent to real sources. Non-individual HRTF-measurements did not only show an overall reduced localization performance, but were particularly prone to front-back-confusions [4], [9], [10]. Møller and colleagues reported about more errors located in the median plane for dummy head HRTFs [11]. Additionally, Andéol and colleagues observed a higher rate of front-back-confusions also for individual HRTFs, but stated greater errors in elevation for the non-individual reproduction via headphones [12].

Dynamic reproduction, allowing for head movements, significantly improved spatial localization in terms of a reduced reversal error [9], [13]–[16]. Indeed, head movements seemed to play an essential role for a realistic sound perception and have been subject to a wide range of studies (e.g. [6], [17]–[19]).

Most of the published work investigating the benefit of dynamic reproduction in combination with individually measured HRTFs suffer from two drawbacks. Either the resolution of the individual HRTFs is rather low [8], [20], [21], which would artificially inflate the measured localization errors, or the tested positions are all located in the horizontal plane [8] which would reduce the expected gain from using individual HRTFs in the first place. Sandvad indeed looked at the influence of HRTF resolution, but used interpolation from relatively low resolution to achieve high resolution samples [20]. An extensive measurement of individual HRTFs was done in [9] but the authors voice concerns about the measurement quality themselfs. Indeed, only the recent development of measurement systems that are capable of acquiring high resolution individual HRTFs in a short period of time (e.g [22]) enable a more thorough investigation.

Similar to Begault and colleagues [8], the present investigation combined and compared individual and non-individual binaural stimuli used in static and dynamic reproduction. Additionally, a minimum resolution necessary for dynamic reproduction was to be estimated, since expenditure of time constitutes a relevant factor for feasibility of individual HRTF-measurements. To cover the range of nearly all audible frequencies, broadband noise pulses were used instead of speech stimuli [8].

Based on previous findings (e.g. [10]), individual HRTFs were expected to provide better results for localization ability. The same holds for dynamic sound reproduction, assumed to be superior to static sound reproduction (e.g. [4]) with an increased localization performance for high HRTF-resolutions.

## 3 Methods and equipment

This chapter will give an overview of all used systems and algorithms.

### 3.1 Binaural measurements and equalization method

One of the aims of the experiment was to determine the influence of the HRTF resolution of dynamic reproduction on the localization accuracy. To this end, three different HRTF resolutions were needed: 5°, 2.5° and 1° in azimuth and elevation, respectively (see section 4).

#### 3.1.1 HRTF measurement

The HRTF measurements took place in a semi-anechoic chamber (*l* × *w* × *h* = 9.2 × 6.2 × 5.0 m^3^) with a lower boundary frequency limit of 200 Hz.

The measurement system was specifically developed for fast individual HRTF measurements with a small impact on the measurement signal [22]. It used 64 loud-speakers (1 inch) arranged in an incomplete semi-circle from elevation 0 ° to 160 ° with 2.5 ° degree resolution (see Figure 1). The participants had two Sennheiser KE3 microphones inserted into their ears at the entrance of the blocked ear canals. They were positioned standing at two meters ear height in the center of the measurement arc on a turntable. The measurement signal was an interleaved sweep of all 64 loudspeakers with a frequency range from 500 Hz to 22, 050 Hz with a time delay between loudspeaker starts of 40 ms and an overall length of 3.38 s.

**Figure 1:**
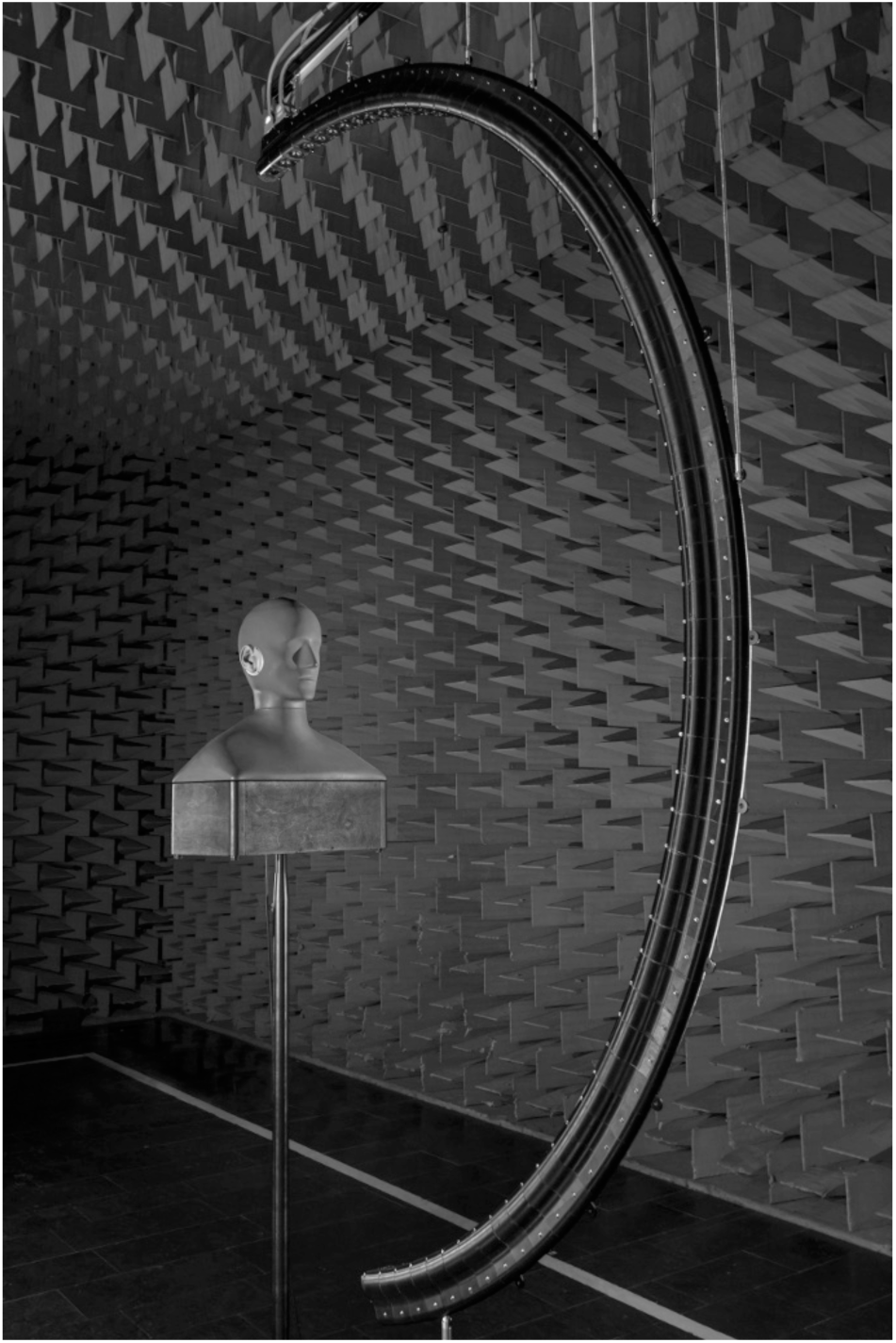
The HRTF Arc with 64 loudspeakers, exemplarily an artificial head is positioned in the center [2].

To reduce unnecessary head movement, the participant’s head was positioned against a head rest. The turntable moved the participant in azimuth angle of 2.5 °. The overall measurement duration was ten minutes.

The dummy head was measured in the same measurement setup with identical measurement signal.

While the 5° resolution could be achieved by simply only using every second azimuth step of the 2.5° measurement data, an interpolation was needed to achieve the 1° resolution.

#### 3.1.2 Interpolation

As shown by Duraiswaini and colleagues as well as Richter and colleagues [23], [24], the spherical harmonic decomposition could be used to interpolate HRTF datasets. To this end, each frequency bin from all measured directions could be regarded as a frequency dependent directivity, measured on a spherical surface. This data was then transformed to spherical harmonic coefficients. The interpolation depended on the maximum order chosen for this transformation. The lower the maximum order was chosen, the more interpolation would occur. It should be noted that this interpolation would not preserve the measured data points as they were also affected.

For this publication, a maximum order of 50 was chosen. As the measured data was not acquired on a whole sphere, regularization as presented by Duraiswaini and colleagues [23] with the regularization factor of 10^−5^ was used to reduce unwanted artifacts. After the decomposition into a finite number of coefficients a reconstruction to arbitrary points could be done. These points were chosen with the same elevation values as the measurement but with a higher azimuth resolution of 1° instead of 2.5°.

#### 3.1.3 HpTF measurement

Headphone-transfer-functions (HpTFs) were measured to calculate an adequate robust equalization. After each of in total eight HpTF measurements, headphones were repositioned on the participants head [25]. To give the best comfort, the repositioning was done by the participant himself/herself. Based on Masiero and Fels [25] the equalization was calculated using the mean of the HpTF measurements. Since phase information was lost at this process, minimum phase was used. Furthermore, notches in the high frequency range were smoothed [25], [26].

### 3.2 Stimuli

Participants were asked to localize a train of pulsed white Gaussian noise. The frequency domain ranged from 100 Hz to 20 kHz providing high-frequency pinna cues essential for localization [27]. The use of white noise was in accordance with previous studies and strongly recommended for the investigations of localization with head movements as broadband signals will excite all changes in monaural in binaural information during tiny head motion (e.g. [16], [28]–[30]).

Head movements were hardly witnessed in case of short sound events smaller than 200 ms, but best observable for minimum duration of 2 s [31]. For this purpose, the total stimulus length was set to 3.7 s and was composed of five alternating 0.3 s- and 1.2 s-bursts of constant sound level with on- and offset ramps (50 ms rise and 50 ms fall time each), interrupted by small pauses of 100 ms.

### 3.3 Real-time binaural synthesis

The stimuli had to be convolved with both HpTF and HRTF filters. As participants were free to move their head, different HRTF filters had to be used, depending on the current head orientation. These filter changes lead to the requirement of a real-time controlled playback that could adjust the playback with no noticeable delay. To this end, a real-time binaural synthesis application and a tracking system to monitor head movement was needed.

#### 3.3.1 Auralization

The real-time auralization was realized with the software Virtual Acoustics (VA) which was developed at the Institute of Technical Acoustics [32]. It allowed for fast exchange of filters as a result of head movement and could be controlled by various tracking systems. It used uniform partitioned convolution [33] and a block size of 128 samples with sampling rate of 44100 Hz. Neighboring HRTFs were selected on a nearest neighbor basis. The latency of the auralization software is in the worst case two block sizes. One internal buffer for the convolution and one hardware buffer of the soundcard. This would calculate to a worst case latency of approximately 6 ms.

#### 3.3.2 Tracking

An optical tracking system was used to monitor several parts of the experiment. The system used four infrared cameras mounted in the room. The system operated with a sampling frequency of 120 Hz and was capable of monitoring several tracking bodies simultaneously. The first monitored motion was the participants head movement. To this end, a tracking body was fastened centrally on the headphone. This point was be monitored throughout the experiment. As this tracking body is not located in the center of the head, the movement of this tracking body did not correspond in all axes to the actual head movement. To correct this offset, a head calibration was done individually. For this calibration, the position of the two ears was marked at the beginning of the test with two additional tracking bodies. From these two positions and the position on top of the head, the center point of the head could be approximated more closely. This movement data was given directly to the auralization software to update HRTF filters if needed. The second use of the tracker was its implementation into the pointing method described in detail in Section 3.4. It involved a marker to indicate spatial positions and an input device, both of which were equipped with tracking markers to monitor the two devices. The latency of the tracker is given by the manufacturer as approximately 8 ms. Combined with the software latency the overall system latency is at worst 14 ms.

### 3.4 Pointing Method

Any investigation into localization accuracy has to be viewed in the context of the used pointing method as the method can heavily influence the achievable accuracy. Over the years, many different approaches to pointing have been proposed and evaluated. Notable methods can be divided into exocentric methods, in which the participant indicates the auditory event on a sphere representing auditory space (e.g GELP (God’s eye view localization point) [34], 2D/3D interfaces [8], [35]) and egocentric methods, in which the participants own center is used as a reference point and pointing is done with an extension of the body (e.g laser pointing [36], [37] or head pointing [21], [38], [39]).

Egocentric methods are generally regarded as more precise [21], [40], [41] and head pointing methods in particular are recommended by [21]. For closed-loop localization however, in which the sound plays during the pointing task, this method will always reflect localization accuracy in the front, as the participant will move their head towards the source.

In this publication, proximal pointing is used as a pointing method. The egocentric method was introduced by Bahu and colleagues [42] and is recommended for closed-loop localization tasks. The listener indicates spatial positions by placing a hand-held marker in the region of the head where (s)he perceives the incident sound (see Figure 2). For this technique the spatial co-ordinate system of the pointing mechanism is concentric with the coordinate system of the stimulus, since localization is referred to the center of the head, or more precisely, to the midpoint of the interaural axis. Closed-loop localization is enabled, as the method does not require head or body rotation. It is practical for both stimuli presentations over headphones and loudspeakers. In this pointing method, only auditory and proprioceptive modalities are involved, neglecting potential visual cues if the hand is positioned in the field of view. According to Bahu and colleagues, proximal pointing yielded more accurate results for high elevations than head and manual pointing. Fast response times may prove advantageous with regard to fatigue and motivation.

**Figure 2:**
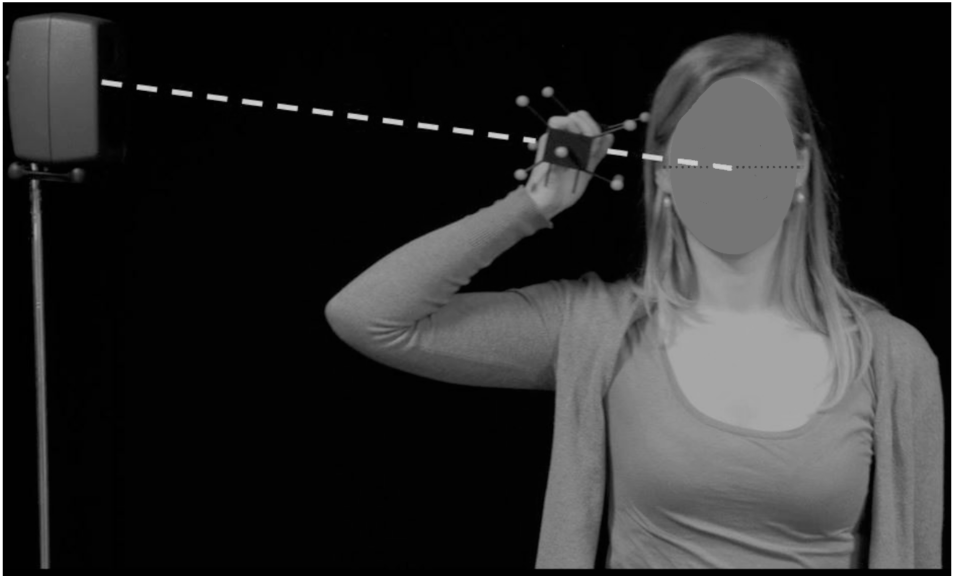
Participant localizing with proximal pointing method.

Localization of sound sources by proximal pointing is a matter of aligning three points of reference: The center of the pointer is positioned onto the connecting line between sound source and the center of the head. To indicate the direction of the perceived auditory stimulus, only the center of the pointer is relevant, its orientation is of no further consequence. Participants were asked to always hold the pointer in the right hand. This limitation ensures an unchanged localization precision without effects due to handedness. As a consequence, source positions were limited to the right hemisphere. To confirm the indicated sound source position, participants were asked to push a button on a second device held in the left hand. The position of this input device was also observed. To indicate an in-head-localization (compare Section 4.2.4) participants were asked to raise their left hand holding the input device from default position up to the height of their head.

### 3.5 Participants

11 right-handed, student participants aged between 20 and 27 years with an average of (23.6 ± 2.4 years) completed the localization experiment. Participants were equally divided in male and female listeners and the experiment was remunerated with 40 Euro. All participants were inexperienced in hearing with HRTF-data and not trained in localization prior to the experiment, however they all achieved a localization accuracy of < 30 ° in the training session when listening to real sources (see Section 3.7.1). Hearing sensitivity (< 15 dB HL) has been verified with the aid of a pure tone audiometry between 125 Hz and 10 kHz.

### 3.6 Room setup

The listening tests took place in a hearing booth (*l* × *w* × *h* = 2.3 × 2.3 × 1.98 m^3^) to ensure a quiet environment during the test. Lights were turned off during the listening test to direct the focus to the aural sense other than the visual sense [43].

### 3.7 Experimental schedule

The experiment was split up in six sessions of about 45 min. each for every participant. While in session 1 individual HRTFs and HpTFs were measured (see Section 3.1) and audiometric data was collected, session 2, 3 and 4 were dedicated to training. Data used for analysis were only taken from session 5 and 6, which were performed within three days. 2 HRTFs × 4 reproduction methods × 11 positions × 4 repetitions + 48 getting-warm trials = 400 trials were collected for data analysis.

#### 3.7.1 Training

To get used to the proximal pointing method, three training sessions with optical feedback on a screen were performed. To be able to give feedback regarding the localization of the source and avoid learning how to localize with a specific HRTF set the training was done with real sources (loudspeakers: Genelec 6010). The training sessions were held in a highly damped room (*l* × *w* × *h* = 8 × 5 × 2.65 m^3^, reverberation time: 0.15 s) with real sources positioned in the right hemisphere of the participant. These source positions were different from the positions used in the listening test itself. Split up into blocks of 25 trials each in total 600 trials were delivered in the three training sessions.

## 4 Experimental design

### 4.1 Independent factors

The independent variables were HRTF (HRTF: individual vs. non-individual), reproduction method (RepMeth: static vs. dynamic (5°) vs. dynamic (2.5°) vs. dynamic (1°)) and position (Pos: Front vs. Side vs. Back) as within-subject variables.

#### 4.1.1 HRTF

HRTFs used in the localization experiment were either individual or measured with an artificial head and therefore non-individual (see Section 3.1 for more information).

#### 4.1.2 Reproduction Method

A static reproduction (i.e. head movements of the participant do not cause any change in binaural reproduction; the same HRTF is used throughout the presentation of the stimulus) was compared to a dynamic reproduction (i.e. head movements of the participant cause a change in binaural reproduction from a certain angle of movement; HRTFs are adjusted throughout the presentation of the stimulus). Three different resolutions of dynamic reproductions were compared: 5°, 2.5° and 1° (see Section 3.1 for more information).

#### 4.1.3 Source position

The factor source position had three levels, categorized from eleven positions scattered in the right hemisphere, all with a distance of 1.5 m from the participant’s center of the head (see Table 1 and Figure 3). There were four frontal positions, two on the median plane with difference in elevation of 30 ° as well as two positioned close to the median plane (*φ* = 350 °). Sources in back were designed to be symmetrically identical with those in front. However, the source with the lower elevation (*θ* = 120 °) was not added due to limitations of the pointing method (see Section 3.4). Sources to the side were also designed to be symmetrically identical and to be positioned on one cone of confusion.

**Table 1:** Angles of the virtual sound source positions in localization experiment, where *φ* indicates the horizontal angle (*φ* = 0 ° indicates the front and increased counterclockwise) and *θ* the vertical angle (*θ* = 0 ° was above the participant and *θ* = 180 ° was underneath the participant.)

| position | $\varphi$ | $\vartheta$ |
| --- | --- | --- |
| front | $0^\circ$ | $60^\circ$ |
| | $0^\circ$ | $90^\circ$ |
| | $350^\circ$ | $90^\circ$ |
| | $350^\circ$ | $120^\circ$ |
| side | $300^\circ$ | $90^\circ$ |
| | $292.84^\circ$ | $70^\circ$ |
| | $247.16^\circ$ | $70^\circ$ |
| | $240^\circ$ | $90^\circ$ |
| back | $190^\circ$ | $90^\circ$ |
| | $180^\circ$ | $60^\circ$ |
| | $180^\circ$ | $90^\circ$ |

**Figure 3:**
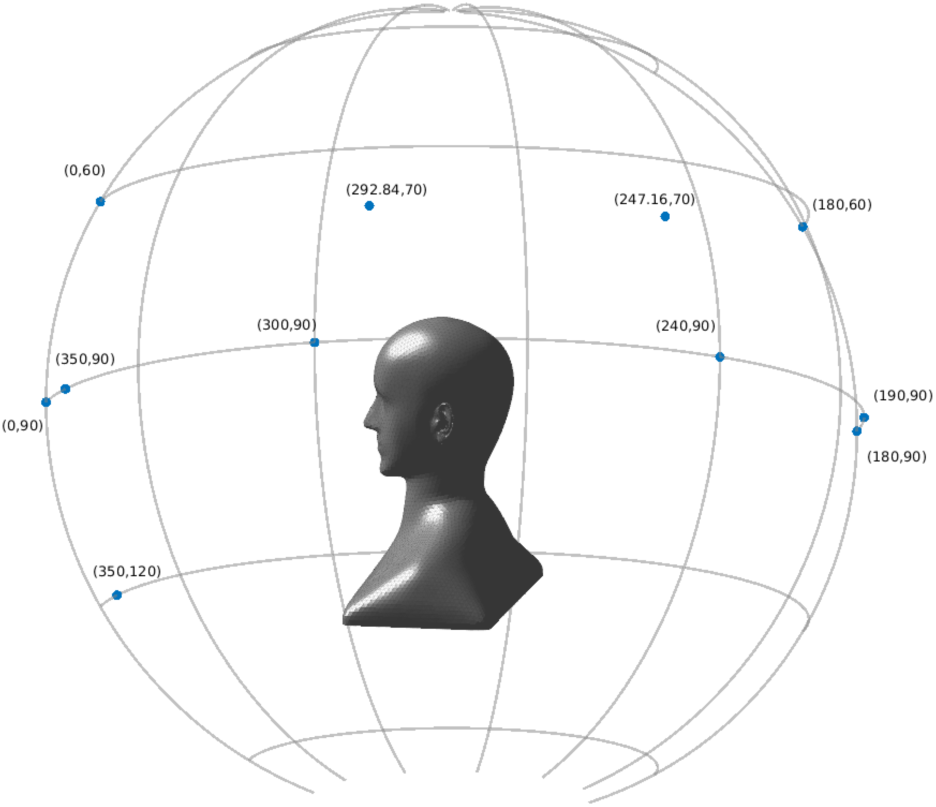
Source positions around listener in localization experiment.

### 4.2 Dependent factors

The dependent variables were localization error and in-head-localization. For a more detailed analysis the localization error was split into three different ways of counting the localization error: the unsigned horizontal error, unsigned vertical error and weighted front-back-confusions.

#### 4.2.1 Unsigned horizontal error

The unsigned error gave a measure of absolute deviation from the correct direction of sound incidence. Deviations from the correct position were added up according to:

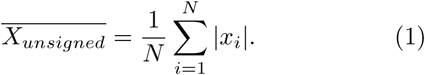

Therefore, the unsigned horizontal error described the absolute deviation of the localized sound position to the actual sound source position in azimuth.

#### 4.2.2 Unsigned vertical error

Accordingly, the unsigned vertical error described the absolute deviation of the localized sound position to the actual sound source position in elevation.

#### 4.2.3 Weighted front-back-confusions

Front-back-confusions as one of the “most important impediment[s] for success of the binaural technique” [44] were taken account of by calculation of the rate of front-back-confusions. A front-back-confusion occurred when the actual sound source was positioned in the frontal hemisphere [270°; 90°), but localized in the rearward hemisphere [90°; 270°). A back-front-confusion described the inverted effect. In the following, there was no distinction drawn between back-front-confusions and front-back-confusions and both were called front-back-confusions. The occurrence of a reversal in a specific trial was denoted with the Boolean

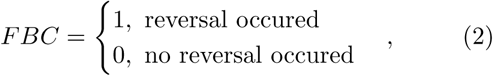

Since front-back-confusions easily occur for lateral source positions, but simply due to localization impression (e.g. actual source position is at 260° and is localized at 275°), a weighting factor should find a remedy:

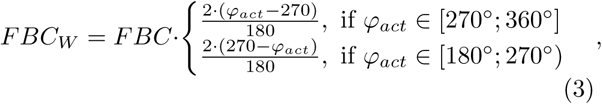

The definition was defined to the right hemisphere where *φ*_*act*_ was the actual sound source position.

#### 4.2.4 In-head-localization

An “in-head-localization” was present when the perceived distance was smaller than the radius of the head [43]. The rate of in-head-localization, was computed thanks to position data of the input device held in left hand indicating a sound perception in the head by raising the arm (see also Section 3.4).

## 5 Results

The presented results were based on data from eleven participants.

In case Mauchly’s test indicated that the assumption of sphericity was violated, the Huynh-Feldt correction was applied. However, for a better overview uncorrected degrees of freedom were reported.

### 5.1 Unsigned horizontal error

Figure 4 shows unsigned horizontal error as a function of the position, reproduction method and the HRTF.

**Figure 4:**
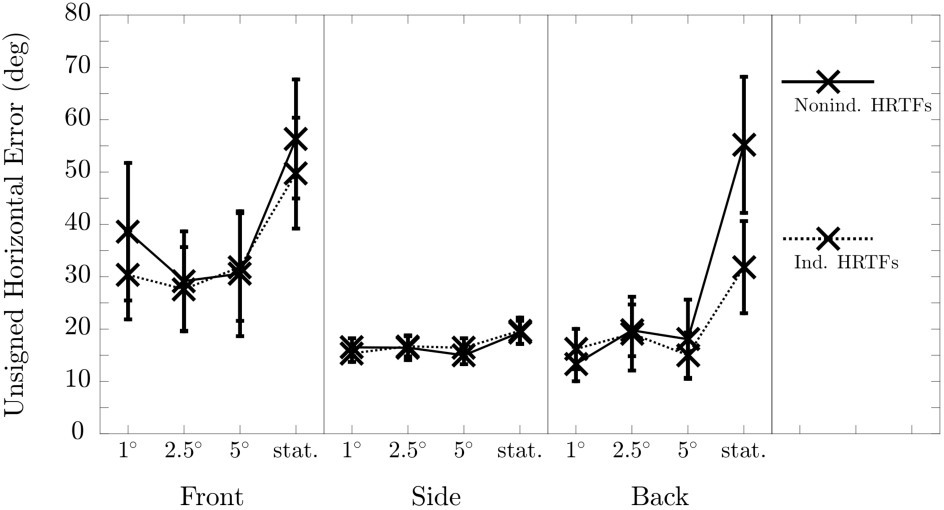
Unsigned horizontal error (in deg) as a function of HRTF, reproduction method and source position (**HRTF** × **RepMeth** × **Pos**). Error bars indicate standard errors.

Although there was a non-significant trend towards smaller horizontal errors when listening to individual HRTFs (individual HRTFs: 24.2 ° vs. non-individual HRTFs: 27.4 °), there was no significant effect on HRTFs in the unsigned horizontal error.

The main effect of reproduction method was significant, resulting in a significantly higher unsigned horizontal error for a static reproduction than for all three dynamic reproduction methods (static: 38.7 ° vs. dynamic (5°): 21.2 ° vs. dynamic (2.5°): 21.5 ° vs. dynamic (1°): 21.7 °). The interaction of HRTF and reproduction method was significant. Post-hoc test showed that the unsigned horizontal error difference between individual and non-individual HRTFs was significantly greater for the static reproduction than for the dynamic reproductions (static: 9.86 ° vs. dynamic (5°): 0.20 ° vs. dynamic (2.5°): 0.64 ° vs. dynamic (1°): 2.18 °).

The main effect of source position was not significant. However, unsigned horizontal errors were smallest for trials where the source was positioned to the side and greatest for trials where the source was positioned in front (Front: 36.8 ° vs. Side: 16.9 ° vs. Back: 23.6 °). As the error was not corrected for front-back confusions this result is not surprising. The interaction of HRTF and source position did not turn out to be significant, meaning that there was gain from the use of individual HRTFs at specific locations only. The interaction of reproduction method and source position was significant, indicating no significant differences between reproduction methods for source positions to the side, but several significant differences between the static and dynamic reproduction for source positions in front and in back. This is explained by a reduction of the number of front-back confusions with dynamic playback.

The three-way interaction was not significant. All p and F-Values for the analysis can be found in Table 2.

**Table 2:** F and p-Values taken from the ANOVA regarding unsigned horizontal error

|  |  |  |
| --- | --- | --- |
| HRTF: | $F(1, 10) = 2.50,$ | $p > .05$ |
| RepMeth: | $F(3, 30) = 15.85,$ | $p < .001$ |
| HRTF × RepMeth: | $F(3, 30) = 3.70,$ | $p < .05$ |
| Pos: | $F(2, 20) = 2.57,$ | $p > .05$ |
| HRTF × Pos: | $F < 1$ | |
| RepMeth × Pos: | $F(6, 60) = 3.48,$ | $p < .05$ |
| HRTF × RepMeth × Pos: | $F(6, 60) = 2.18,$ | $p > .05$ |

### 5.2 Unsigned vertical error

Figure 5 shows unsigned vertical error as a function of the position, reproduction method and the HRTF. There was no significant effect in the unsigned vertical error for either the used HRTF, the reproduction method or the interaction between both. This indicates, that there was no gain in the elevation localization accuracy with the use of individual HRTFs regardless of reproduction method. The main effect of source position was significant, indicating significantly smaller unsigned vertical errors for trials where the source was positioned to the side than for trials where the source was positioned in front or in back (Front: 19.2 ° vs. Side: 10.3 ° vs. Back: 20.6 °). The two-way interactions with source position as well as the three-way interaction did not turn out to be significant. All p and F-Values for the analysis can be found in Table 3.

**Table 3:**
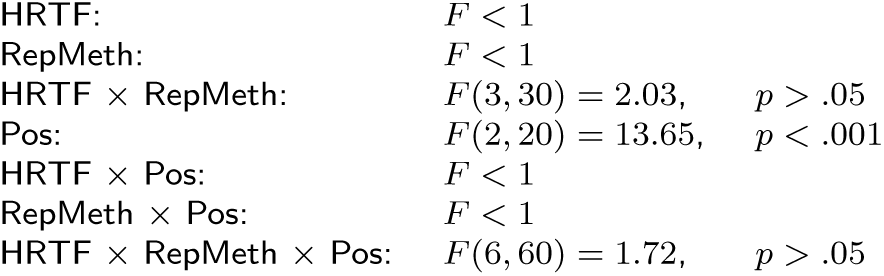
F and p-Values taken from the ANOVA regarding unsigned vertical error

|  |  |  |
| --- | --- | --- |
| HRTF: | $F < 1$ | |
| RepMeth: | $F < 1$ | |
| HRTF × RepMeth: | $F(3, 30) = 2.03,$ | $p > .05$ |
| Pos: | $F(2, 20) = 13.65,$ | $p < .001$ |
| HRTF × Pos: | $F < 1$ | |
| RepMeth × Pos: | $F < 1$ | |
| HRTF × RepMeth × Pos: | $F(6, 60) = 1.72,$ | $p > .05$ |

**Figure 5:**
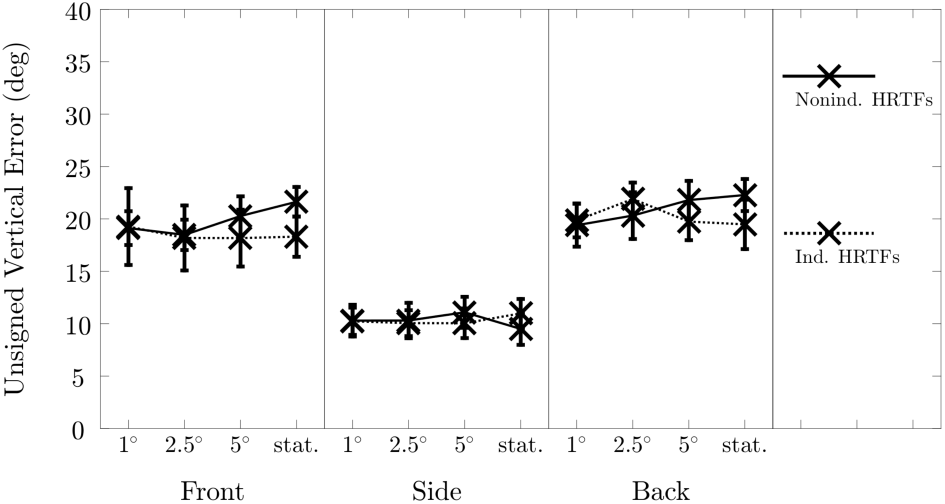
Unsigned vertical error (in deg) as a function of HRTF, reproduction method and source position (**HRTF** × **RepMeth** × **Pos**). Error bars indicate standard errors.

### 5.3 Weighted front-back-confusions

Figure 6 shows weighted front-back confusion in percent as a function of the position, reproduction method and the HRTF. The arc-sine transformation of data which is appropriate for the data on proportions is applied before conducting the ANOVA. The ANOVA yielded a significant effect on HRTFs in the weighted front-back-confusions, indicating less front-back-confusions when listening to individual HRTFs (individual HRTFs: 7.6 % vs. non-individual HRTFs: 10.7 %).

**Figure 6:**
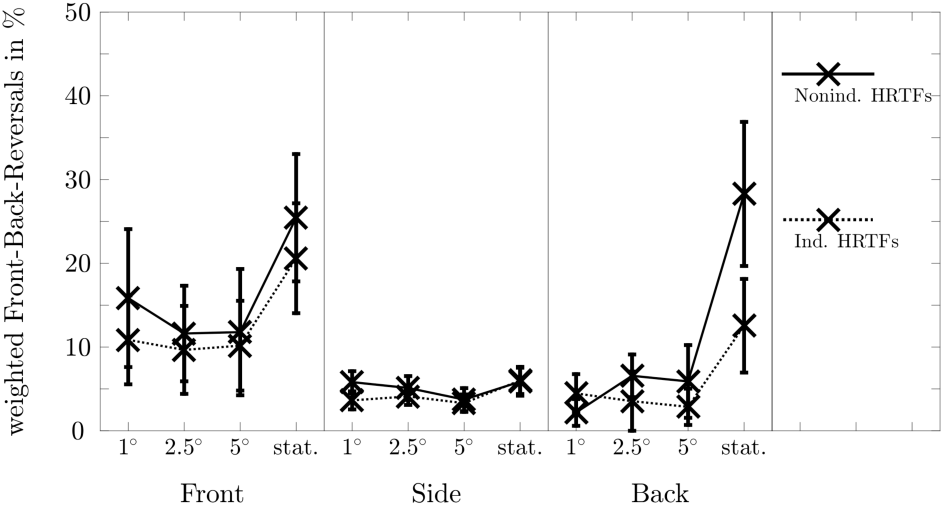
Weighted front-back-confusions (in %) as a function of HRTF, reproduction method and source position (**HRTF** × **RepMeth** × **Pos**). Error bars indicate standard errors.

**Figure 7:**
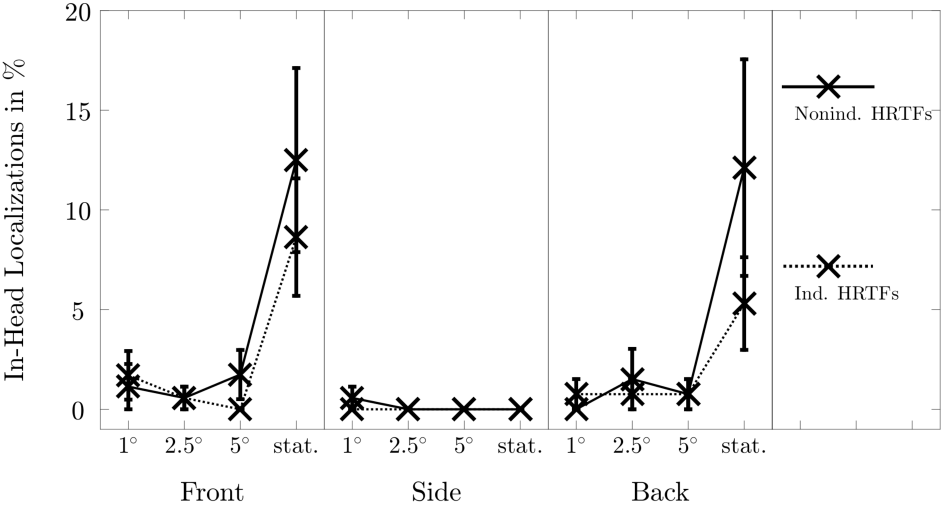
In-head-localization (in %) as a function of HRTF, reproduction method and source position (**HRTF** × **RepMeth** × **Pos**). Error bars indicate standard errors.

The main effect of reproduction method was also significant, resulting in a significantly higher front-back-confusion for a static reproduction than for all three dynamic reproduction methods (static: 16.5 % vs. dynamic (5°): 6.3 % vs. dynamic (2.5°): 6.8 % vs. dynamic (1°): 7.1 %). The interaction of HRTF and reproduction method was not significant. However, there was a non-significant trend towards a greater front-back-confusion difference between individual and non-individual HRTFs for the static reproduction than for the dynamic reproductions (static: 6.8 % vs. dynamic (5°): 1.7 % vs. dynamic (2.5°): 2.0 % vs. dynamic (1°): 1.6 %). This leads to the interpretation that dynamic reproduction will reduce the gain from individual HRTFs in terms of front-back confusion.

The main effect of source position was not significant. However, weighted front-back-confusions were smallest for trials where the source was positioned to the side and greatest for trials where the source was positioned in front (Front: 14.5 % vs. Side: 4.7 % vs. Back: 8.3 %). The two-way interaction of reproduction method and source position is significant, indicating higher front-back-confusions for a static reproduction in front and in back compared to the side. The two-way interaction of HRTF and source position as well as the three-way interaction did not turn out to be significant. All p and F-Values for the analysis can be found in Table 4.

**Table 4:** F and p-Values taken from the ANOVA regarding weighted front-back-confusions

|  |  |  |
| --- | --- | --- |
| HRTF: | $F(1, 10) = 6.98,$ | $p < .05$ |
| RepMeth: | $F(3, 30) = 11.68,$ | $p < .001$ |
| HRTF × RepMeth: | $F(3, 30) = 1.31,$ | $p > .05$ |
| Pos: | $F(2, 20) = 1.31,$ | $p > .05$ |
| HRTF × Pos: | $F(2, 20) = 1.02,$ | $p > .05$ |
| RepMeth × Pos: | $F(6, 60) = 3.54,$ | $p < .05$ |
| HRTF × RepMeth × Pos: | $F(6, 60) = 2.17,$ | $p > .05$ |

### 5.4 In-head-localization

Figure 6 shows in-head localization occurance in percent as a function of the position, reproduction method and the HRTF. The arc-sine transformation of data which is appropriate for the data on proportions is applied before conducting the ANOVA. There was no significant effect on HRTFs in in-head-localization. However, there was a non-significant trend towards a lower rate of in-head-localization when listening to individual HRTFs (individual HRTFs: 1.5 % vs. non-individual HRTFs: 2.6 %).

The main effect of reproduction method was significant, resulting in a greater rate of in-head-localization for the static reproduction than for all three dynamic reproduction methods (static: 6.4 % vs. dynamic (5°): 0.5 % vs. dynamic (2.5°): 0.6 % vs. dynamic (1°): 0.7 %). The interaction of HRTF and reproduction method was not significant. However, there was a non-significant trend towards a greater in-head-localization difference between individual and non-individual HRTFs for the static reproduction than for the dynamic reproductions (static: 3.6 % vs. dynamic (5°): 0.5 % vs. dynamic (2.5°): 0.3 % vs. dynamic (1°): −0.2 %).

The main effect of source position was significant, indicating a significantly lower rate of in-head-localization for trials where the source was positioned to the side than for trials where the source was positioned in front (Front: 3.4 % vs. Side: 0.1 % vs. Back: 2.7 %). The two-way interaction of HRTF and source position did not turn out to be significant. The interaction of reproduction method and source position was significant, indicating more in-head-localization with the static reproduction for sources positioned in front and back than to the side.

The three-way interaction was not significant. All p and F-Values for the analysis can be found in Table 5.

**Table 5:** F and p-Values taken from the ANOVA regarding in-head-localization

|  |  |  |
| --- | --- | --- |
| HRTF: | $F(1, 10) = 1.92,$ | $p > .05$ |
| RepMeth: | $F(3, 30) = 8.12,$ | $p < .05$ |
| HRTF × RepMeth: | $F(3, 30) = 1.68,$ | $p > .05$ |
| Pos: | $F(2, 20) = 6.59,$ | $p < .05$ |
| HRTF × Pos: | $F < 1$ | |
| RepMeth × Pos: | $F(6, 60) = 5.36,$ | $p < .05$ |
| HRTF × RepMeth × Pos: | $F(6, 60) = 1.63,$ | $p > .05$ |

## 6 Discussion

In general, lateral sound sources yielded greater localization precision compared to those in front and back (e.g. [3]). This assumption was validated by the present investigation. There was a precision significantly higher by about 10° in elevation for lateral positions compared to frontal and backward positions and almost no and therefore a significantly lower rate of in-head-localization were found at the side. Front-back-confusion rates tended to be less pronounced for positions at the side and a tendency towards generally higher localization accuracy in azimuth for lateral positions was found in the present data.

One of the main investigations of this publication was the influence of the resolution of the HRTF. The hypothesis was that a higher resolution HRTF will increase the localization performance during dynamic reproduction. This hypothesis could not be confirmed. There were no identifiable differences between the different resolutions. Sandvad, conducting listening experiments with interpolated HRTFs with resolution of 1 ° up to 13 °, reported similar findings: “Spatial resolution appears to have very little influence on subject performance. Large angular steps between different HRTF filters introduces audible discontinuities in the sound, but as with reduced update rate, participants seemed to be able to ignore the audible artifacts, and complete the task with good precision [20].” These artifacts could also be observed: Several participants stated after the present experiment that some trials suffered from audible discontinuities in the sound. Due to the design of the experiment there was no chance to keep track of the specific affected trials. However, the authors themselves also perceived these audible discontinuities in sound during test runs and they could all be reduced to the dynamic reproduction with a resolution of 5 °. Thus, a resolution smaller than 5 ° should be recommended, even though there is no increase in localization performance.

The results of this investigation could confirm that dynamic reproduction enables a more accurate localization compared to static reproduction. Dynamic reproduction was observed to reduce the unsigned horizontal error by 17° and reversal rate could be lowered by more than 9 %. This was in accordance with Begault and colleagues who measured confusion rate differences up to 31 %, stating: “Head tracking reduced reversals significantly (front-back- and back-front-confusions of the location of the stimuli across the inter-aural axis from a rate of 59 % to 28 %)[13].” Taking account of front-back-confusions in the unsigned horizontal error by extracting the affected trials, no effect is discernible (Unsigned horizontal error corrected by discarding all front-back-confusion trials: [RepMeth: *F* < 1]). Therefore, the unsigned horizontal error was strongly correlated with the front-back-confusion rate. As a consequence, dynamic reproduction effectively improved sound localization by eliminating, or, at least, reducing the confusion rate. The very same dynamic cues resulted advantageous also for reduction of in-head-localization rates. As a general result, dynamic reproduction seemed to have a great effect on reduction of typical problems arising with virtual acoustic environments such as front-back-confusion and in-head-localization.

One of the major aims of the presented experiment was to figure out, up to which extent individual HRTFs prove advantageous with respect to non-individual HRTFs. A significant main effect was observed only once: On average, front-back-confusions were reduced by about 3 % when individual HRTFs were used. Letowski and Letowski, noted slightly higher reversal rates ranging around 7-11 % [45]. These findings were not in line with those by Begault and colleagues who did not observe a significant main effect for the HRTF analyzing the front-back-confusion rate [13]. An impact on unsigned horizontal error due to individual HRTFs could be reasonably assumed, however, in the present experiment no significant main effect for the HRTF was observed. The absence of the main effect could be explained by the imbalance of static and dynamic reproduction methods (1:3) and that there is no effect for lateral positions. By reducing the variable of reproduction methods to two levels of static vs. dynamic (5 °) and neglecting lateral positions a marginally significant main effect on HRTFs could be found [HRTF: *F* (1, 10) = 4.86, *p* = 0.052]. Regarding the accuracy of localizing in elevation no difference between individual and non-individual binaural stimuli could be found. In contrast to that, Andéol and colleagues found considerable differences in the low elevation region [11], [12], substantiating the fact that individual binaural stimuli yield a better localization accuracy in elevation than non-individual.

The experiment was designed to observe interactions between different HRTFs and Reproduction Methods, which turned out to be significant only for the unsigned horizontal error. However, a tendency towards greater differences between individual and non-individual HRTFs in static than in dynamic reproduction could be observed for all dependent variables. These findings were in agreement with the observations by Begault and colleagues [13]. Large differences in static vs. dynamic reproduction appeared to mask potential effects of the HRTF. The same finding was confirmed by Kato and colleagues who report “that individual differences in HRTFs can be overcome by head motion” [18].

## 7 Conclusion

Aim of this investigation was to find an appropriate setting for the use of individual/non-individual HRTF and static/dynamic reproduction methods to ensure a most realistic auralization of acoustic scenes with binaural technique and headphone reproduction.

Dynamic reproduction of any resolution applied was confirmed fundamental for a reduction of undesired front-back-confusions and in-head-localization. With front-back-confusions dominating the results for the unsigned horizontal error, dynamic reproduction implicitly also had a great impact on the azimuth error. Individually measured HRTFs showed a smaller effect on localization accuracy compared to the influence of dynamic sound reproduction. They were mainly observed to reduce the front-back-confusion rate. In general, individual HRTFs proved indeed beneficial compared with non-individual HRTFs, however, as the benefit tended to be masked in the presence of the more pronounced effect of reproduction method, using non-individual HRTFs in a dynamic reproduction might be a less complex and elaborate way to successfully provide for a realistic auditory perception. The resolution of the used HRTFs did not produce a significant increase in localization performance, however audible artifacts with a resolution of (5 °) were present which signified the need for a higher resolution.

